# Fisetin-mediated MYC restoration improves age-associated decline in macrophage function

**DOI:** 10.64898/2026.06.10.731325

**Authors:** Joshua V. Kimble, Charlotte E. Moss, Lilia S. Hodge, Martha L. Clements, Alexandra Rayson, Kellie S Roberts, Ryan J.H. West, Iwan R. Evans, Sheila E. Francis, Ilaria Bellantuono, Endre Kiss-Toth, Heather L. Wilson

## Abstract

Immune decline in older adults is associated with increased susceptibility to infection and chronic inflammatory diseases. Macrophages are critical innate immune cells that show reduced capacity for phagocytosis and migration with age. Our previous work shows that reduced levels of MYC and USF1 transcription factors are drivers of macrophage age-related functional decline. Here we show that macrophage-specific *Myc* overexpression improves macrophage migration and, more importantly, is able to improve physical performance at older age in *Drosophila*, while lifespan remains unaffected. Treatment of human primary macrophages from older individuals with the geroprotective supplement fisetin reverses the decline in *MYC* expression and improves phagocytosis of pathogens and cell migration functions towards levels seen in younger individuals. Mechanistically, fisetin acts via MYC, by restoring expression levels of MYC targets in human macrophages that are altered with age. Finally, fisetin feeding in older mice improves motor activity and reduces frailty, as well as restoring primary macrophage function and *Myc* expression *in vitro*. These findings reveal that restoration of MYC in macrophage ageing is responsible, at least in part, for improvement in physical performance with age and identify this pathway as a rational target to reverse age-related immune decline.

**Graphical Abstract:** 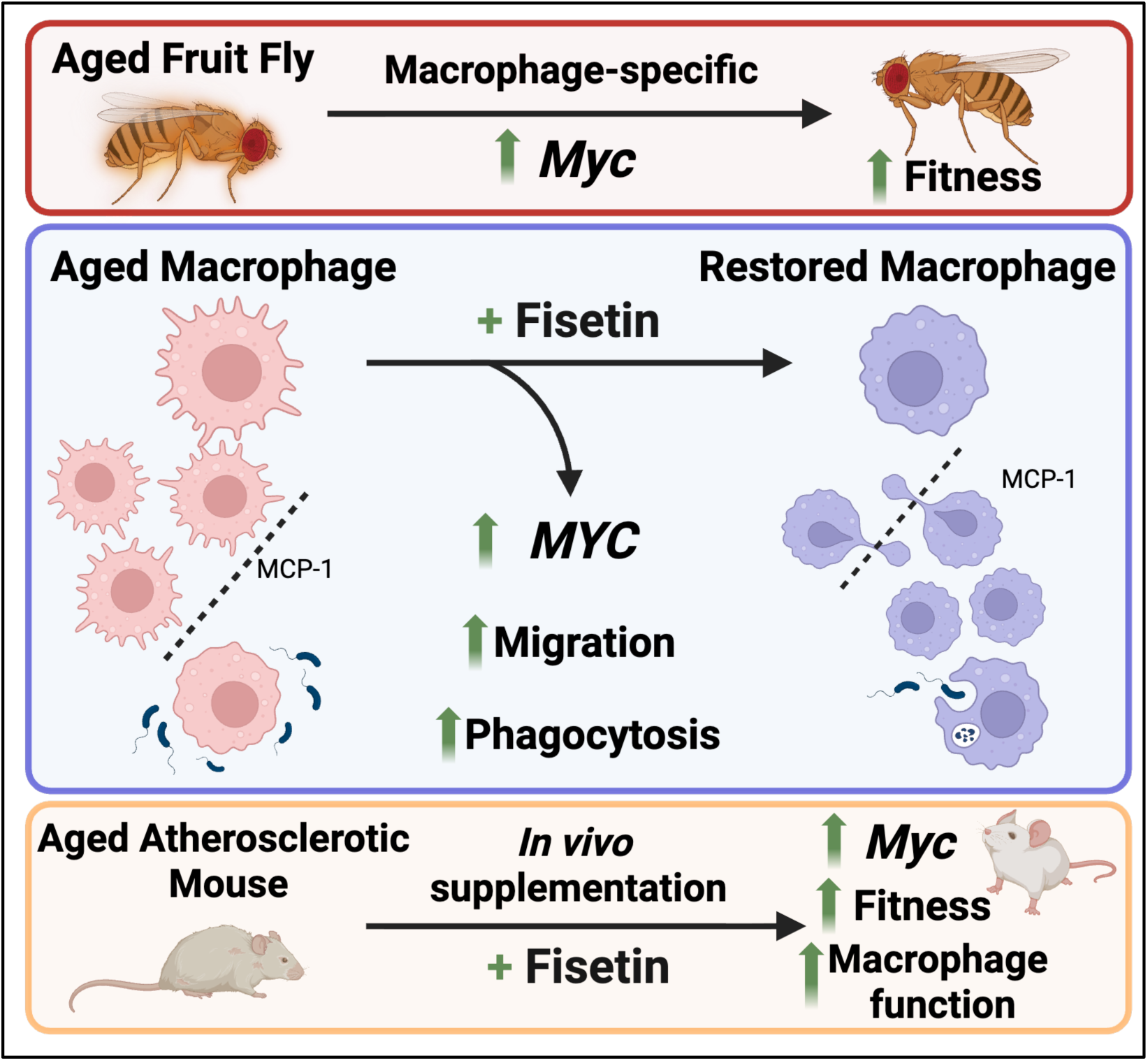

## Introduction

The global prevalence of multimorbidity is growing due to changes in lifestyle risk factors and longer life-expectancy as a result of improved survival from acute and chronic diseases^1^. Ageing is associated with a progressive decline in immune function, leading to increased susceptibility to infection and to chronic inflammatory diseases^2^. Immunosenescence, a decline in immune system function affecting both innate and adaptive immunity, reduces the capacity to respond to infections, vaccination and maintain tissue homeostasis^3^. Age-related immune dysfunction is further characterised by inflammaging, a state of chronic, low-grade systemic inflammation from persistent activation of innate immune pathways, accumulation of senescent cells, and dysregulated immune responses^4^. Inflammaging represents a shared risk factor and pathophysiological mechanism underlying multiple age-associated chronic diseases including cardiovascular disease, diabetes mellitus, chronic kidney disease, cancer, and neurodegenerative disorders, whilst also contributing to frailty, disability, and premature mortality^5,6^.

Macrophages are key cells in the innate immune response, maintaining tissue homeostasis, controlling infections, and regulating metabolic function^7^. They undergo age-associated functional decline, exhibiting reduced phagocytic capacity and impaired migration to chemotactic signals^8^. The decline in macrophage-mediated clearance of pathogens and cellular debris drive infection susceptibility and accumulation of tissue damage. In the cardiovascular system, impaired efferocytosis promotes atherosclerotic plaque formation and instability^9^, whilst dysfunctional macrophages in metabolic tissues exacerbate chronic inflammation, insulin resistance, and type 2 diabetes^10^. Similarly, impaired macrophage-mediated tissue repair mechanisms contribute to musculoskeletal frailty and delayed recovery from illness^11^. Collectively, these multisystemic macrophage deficits establish a mechanistic link between immunosenescence and age-associated multimorbidity.

We recently identified *MYC* as a major driver of age-associated macrophage dysfunction^8^. Primary macrophages from older humans and mice exhibit reduced *MYC* expression, leading to impaired phagocytosis, chemotaxis and migration capacity^8^. Knockdown of *MYC* in monocyte-derived macrophages (MDMs) from young donors reproduces these age-associated functional deficits, establishing *MYC* as a central regulator of macrophage ageing^8^. These findings suggest that restoring *MYC* expression in aged macrophages could reverse functional decline and enhance innate immune responsiveness, establishing *MYC* and its downstream targets as a potential therapeutic target. However, pharmacological interventions that can restore *MYC* expression in aged hMDMs remain unexplored, as does the impact of *MYC* restoration on health-span and lifespan *in vivo*.

We therefore assessed whether restoring macrophage MYC expression through genetic or pharmacological interventions could reverse age-associated functional decline and extend health-span in fly and mouse models of ageing and explored molecular mechanisms underpinning *in* vivo changes in primary human and mouse macrophages.

Macrophage-specific MYC and USF1 overexpression in *Drosophila* showed that macrophage-specific MYC overexpression improves cell migration and physical performance, while lifespan remains unaffected. Treatment of human primary macrophages from older individuals with the geroprotector fisetin upregulates MYC expression and improves phagocytosis of pathogens and cell migration towards levels seen in macrophages from younger individuals. Mechanistically, fisetin acts at the MYC axis by co-regulating MYC targets in human macrophages that are altered with age. Finally, fisetin feeding in older mice improves motor activity and health span, restoring macrophage function and MYC expression in bone-marrow derived macrophages. Together these findings define MYC as a central, evolutionally conserved regulator and a rational therapeutic target of macrophage health and show fisetin-mediated restoration of MYC expression in cells from older individuals improves immune function and health-related motor activity.

## Methods

### Drosophila husbandry

*Drosophila melanogaster* stocks were maintained at 25°C on standard cornmeal/agar/molasses medium (Supplementary table 4) in the presence or absence of RU486. For conditional overexpression of *Myc* in adult haemocytes (*Drosophlia* blood cells), male flies carrying *w^1118^/Y*;*UAS-Myc.Z* (Bloomington Drosophila Stock Centre [BDSC] line 9674;^12^ were crossed with virgin female flies carrying *w^1118^*;;*Hml(Δ)-GeneSwitch*^13^ in straight-sided bottles to generate F1 offspring with the genotype *w¹¹¹⁸; UAS-Myc.Z/+; Hml(Δ)-GeneSwitch/+*. Flies were tipped to new bottles every 2 days to expand experimental progeny from the same set of parental flies.

For the lifespan and climbing assays, flies of the appropriate sex and genotype were aged on standard fly media containing RU486 (Fisher - product code 459980050; “RU food”; 3.44 ml of 10 mg/ml stock dissolved in absolute ethanol per 400 ml of fly media to give a final concentration of 200 µM RU486) or fly media containing an equivalent volume of ethanol vehicle (“control food”). Food was cooled to 50-55°C before addition of RU/ethanol vehicle before pouring into vials, set overnight then bunged, stored for up to one month. Haemocyte-specific Myc expression via *Hml(Δ)-GeneSwitch* was induced via culture on RU food, since RU486 binds to the progesterone receptor domain within the GeneSwitch protein^14^, activating GAL4-mediated transcription. Specificity of gene expression in the presence of RU486 via the GeneSwitch system within haemocytes was confirmed via culture of *w^1118^;Hml(Δ)-GeneSwitch/UAS-Stinger* (nuclear GFP^15^) embryos on RU or control food. L3 wandering larvae were picked from those vials, washed in distilled water, and imaged in water on ice using a Leica M205FA fluorescent stereomicroscope (ET-GFP filter, 2x PLANAPO objective lens) and LasX imaging software.

### Live imaging of *Drosophila* embryonic macrophages

Stage 15 *Drosophila melanogaster* embryos containing GFP-labelled haemocytes were generated by crossing virgin females containing pan-haemocyte Gal4 drivers and *UAS-GFP* transgenes (*w^1118^;srpHemo-GAL4*, *UAS-GFP;crq-GAL4*, *UAS-GFP* [abbreviated to *srpGFP;crqGFP*]) with either *w^1118^* control males or *w^1118^;UAS-Myc.Z* males to obtain constitutive macrophage-specific overexpression of *Myc* (*Myc* OE). For experiments investigating Usf function, w* P{EP}UsfG717 (BDSC line 33460) virgin females were crossed to srpGFP;crqGFP males to generate constitutive macrophage-specific overexpression of Usf (Usf OE); n.b., EP lines contain UAS sites enabling GAL4-mediated expression from their insertion positions.

Embryos were collected from apple juice agar plates taken from laying cages incubated at 22°C. Embryos were then washed and dechorionated in bleach for 2 minutes before being rinsed thoroughly in dH₂O. Dechorionated embryos were aligned ventral-side-up on glass slides and covered with a small drop of Voltalef oil (Oil 10S; VWR – product code 24627.188) with imaging commencing 30 minutes after addition of oil. Live imaging was performed using an UltraView Spinning Disk confocal system (PerkinElmer) mounted on an inverted microscope with a 40× UplanSApo oil-immersion objective (NA 1.3). Z-stacks spanning 30 µm with 1 µm step size were acquired and maximum intensity projections were generated in Fiji/ImageJ (v2.14.0/1.8.0). Time-lapse movies of haemocytes between the epithelium and ventral nerve cord were recorded every 2 minutes for 20 minutes to quantify plasmatocyte morphology and migration (n.b., only plasmatocytes [the macrophage equivalents in this model] reach this localisation in the embryo; we refer to these cells as macrophages henceforth).

### Analysis of *Drosophila* embryonic macrophage morphology and migration

#### *Drosophila* embryo plasmatocyte migration analysis

Morphology measurements were performed on the first timepoint of the 20-minute time-lapse migration recordings using Fiji. Individual macrophages on the ventral midline were outlined using the polygon selection tool to define either the whole-cell region or the cell-body region; whole-cell area, cell-body area and perimeter were measured automatically for each cell. Lamellipodial area was calculated as: lamellipodial area = whole-cell area − cell-body area.

To extract migratory parameters, time-lapse image series were converted to TIFF format and maximum projections were assembled in Fiji. These were blinded using the Blind Analysis Tools plugin, and global embryo drift was corrected using StackReg before cell tracking. The Manual Tracking plugin was used to track all macrophages visible in the initial field of view by following the centre of the cell body throughout the 20-minute movie with tracks terminated when cells exited the field. Tracking coordinates were exported as .csv files and analysed using the Chemotaxis and Migration Tool (Ibidi) in ImageJ to calculate migration speed and directionality per cell. Directionality was defined as Euclidean distance across the period tracked / total accumulated distance across that time.

Directionality values range from 0 (non-directional) to 1 (linear migration). For each embryo, mean directionality and mean speed across all tracked cells were calculated to provide a single value per embryo for statistical analysis. Ten embryos per genotype were analysed for migration parameters, using one-way ANOVA with Dunnett’s post-hoc test to compare *Myc*OE with control.

#### *Drosophila* larval haemocyte isolation for RNA extraction

Third instar wandering larvae were collected and washed in distilled water to remove food debris before being dried on tissue and transferred to a 100 μl droplet of chilled PBS on a petri dish lid with one larva dissected per 100 μl droplet. Larval haemocytes were released by grasping the posterior spiracles with forceps and gently pulling apart the larval cuticle to open the body cavity, allowing release of haemocytes into the surrounding medium. The larval carcass was shaken for 10 seconds with forceps to release more blood cells and then discarded; the cell suspension containing haemocytes was collected and kept on ice. Haemocytes from 20 larvae of the same genotype were pooled in 1.5 ml Eppendorf tubes and centrifuged at 1000 rpm for 5 minutes to pellet cells. The supernatant was removed, and cell pellets were stored at −80°C until RNA extraction. Three independent biological replicates were prepared from separate cages for each experimental condition, with each replicate consisting of haemocytes pooled from 20 larvae.

#### *Drosophila* lifespan assay

Bottles in which experimental progeny had been laid were cleared and new flies allowed to emerge for 24h. Larvae were then transferred to fresh bottles without selection or exposure to CO_2_ (day 0). After 2 days (day 2), females were selected using CO2-mediated anaesthetisation and 20 mated females transferred to each vial containing control or RU486 media for ageing at 25oC. For survival analysis, 120 flies per replicate across 6 vials per condition were aged across 3 independent replicates (360 flies in total overall). Flies were tipped to a new vial of RU/control food every 2 days to maintain RU486 activity and mortality was scored during these medium changes. Survival curves were analysed using Kaplan-Meier analysis with log-rank tests.

#### *Drosophila* climbing performance assay

As a measure of fitness, climbing performance was assessed using negative geotaxis assays at weekly intervals from 7-63 days post-eclosion. Flies were placed without anaesthetisation in glass boiling tubes on a white, backlit background. After a 30-second acclimatisation period, video recording was started, and the flies were startled by tapping the apparatus to initiate climbing from the bottom of the tube. Recording continued for 20 seconds with videos captured at 30 frames per second using a Basler ace acA2040-90um NIR camera (12 mm HP-series fixed focal length lens; Basler AG) positioned at a fixed distance and processed using ImageJ/Fiji software with the TrackMate plugin for automated tracking. Individual fly movement was tracked between frames using the Hessian detector with object diameter set to 2 mm and Simple LAP tracker with linking max distance of 10mm. Median climbing speed was calculated over the recording period for each individual fly. To control for circadian effects, all assays were performed within the same one-hour timeslot each week. Flies from three independent cross replicates, with 10 flies per replicate, were analysed at each timepoint, giving a starting population of 30 flies per genotype; exact sample sizes at each timepoint declined due to natural mortality and are detailed in Supplementary Table 5. Flies were transferred to fresh food after each weekly timepoint.

### Mice

C57BL/6J male mice (WT) purchased from Charles River Laboratories (UK) were used at 20 months of age for experiments. Mice were acclimatised to the facility for 2 weeks before starting the study. Mice were housed in groups with a 12h:12h light-dark cycle with ad libitum access to drinking water and food. Mice received a single injection of 1011 vg rAAV8/D377Y-mPCSK9 followed, after one week, by a Western diet for 12 weeks to induce atherosclerosis. Mice were culled at 19 months. All animal experiments were approved by the University of Sheffield Ethical Review Committee and performed in accordance with UK Home Office Project Licence P5395C858.

### Mouse fisetin supplementation

Mice were arbitrarily allocated into two groups: group 1 (aged control) were fed a Western diet (Envigo TD. 88137) from the age of 16 months, group 2 (aged + fisetin) were fed a Western diet (Envigo TD.88137) from the age of 16 months incorporating fisetin (Indofine Chemical Co, NJ) at 500 mg/kg^16^, given for 2 weeks followed by a 2-week break, repeated for a total of 12 weeks.

### Mouse bone marrow-derived macrophage isolation and culture

Bone marrow isolation was performed as previously described^8^. Briefly, bone marrow was isolated from femurs and tibias of aged (20 months) male C57BL6 mice under aseptic conditions. Bones were cut at both ends and flushed with DMEM (Gibco) using a syringe and 25-gauge needle into 10 mL of DMEM. This cell suspension was passed through a 40 μm cell strainer and centrifuged at 500 g for 5 minutes. Cells were then resuspended in DMEM containing 10% (v/v) HI-FBS, 10% (v/v) L929-conditioned (M-CSF rich) media and 1% (v/v) penicillin-streptomycin (Gibco) for 5 days at 37 °C and 5% CO_2_. Differentiated bone marrow-derived macrophages were then used for functional assays or were polarised for a further 24 hours towards an inflammatory phenotype using 100 nM lipopolysaccharide (LPS) (Enzo) and 20 nM interferon-γ (IFNγ) (Peprotech) or towards an anti-inflammatory phenotype using 20 nM interleukin-4 (IL-4) (Peprotech), used for gene expression analysis.

### Mouse bone marrow-derived macrophage polarisation

Following 5 days of differentiation in M-CSF rich media, mouse bone marrow-derived macrophages (BMDM) were either left unstimulated (M^0^) or polarised for 24 hours. For pro-inflammatory (M^1^) polarisation, cells were stimulated with 100 ng/mL lipopolysaccharide (LPS; *Escherichia coli*, Enzo Life Sciences) and 20 ng/mL recombinant mouse interferon-γ (IFN-γ; Peprotech) (M^LPS+IFNγ^). For anti-inflammatory (M^2^) polarisation, cells were stimulated with 20 ng/mL recombinant mouse interleukin-4 (IL-4; Peprotech) (M^IL-4^). Following polarisation, cells were used for functional assays or lysed for RNA extraction and gene expression analysis.

### Mouse rotarod

Motor coordination and balance were assessed using an accelerating rotarod apparatus, as previously described^17^. Briefly, mice were trained on the apparatus for three consecutive days prior to testing. On the test day, mice were placed individually on a rotating rod that accelerated from 4 to 40 rpm over a maximum duration of 300 seconds. The latency to fall was recorded for each mouse (in seconds). Mice that fell within the first 10 seconds were repositioned on the rod. If a mouse fell three times within this initial period, the recorded latency was set to zero and the mouse was removed from the test. Each mouse underwent three trials, with a 15-minute inter-trial interval. Performance was calculated as the average latency to fall across the three trials, with longer latencies indicating better neuromuscular coordination and endurance.

### Mouse grid hanging bar

Muscle strength and endurance were assessed using a horizontal grid test, as previously described^17^. Briefly, each mouse was placed on the grid such that it gripped the grid with all four limbs, after which the grid was inverted. The latency to fall was recorded for a maximum duration of 7 minutes. Mice that fell within the first 10 seconds were repositioned on the grid and allowed an additional trial. If a mouse fell twice within this initial period, the latency was recorded as zero. Each mouse underwent three trials, with a minimum rest period of 30 minutes between trials. Performance was calculated as the average hanging time across the three trials.

### Mouse frailty index

A clinical frailty index was calculated based on the assessment of 29 standardized health deficits. Frailty measures were adapted from the protocol described by Whitehead et al.^18^, with the exception of body weight (excluded due to obesity in the mice) and body temperature. Each deficit was scored as 0 (no deficit present), 0.5 (mild or intermediate deficit), or 1 (severe deficit present). The frailty index score for each mouse was calculated as the proportion of deficits present relative to the total number of deficits assessed. Higher frailty index scores indicate greater frailty.

### Human primary monocyte-derived macrophages

Peripheral blood mononuclear cells (PBMCs) were isolated from healthy donors, in accordance with ethics approved by the University of Sheffield ethics committee (031330 and 064649). All blood donors gave informed consent, and studies were performed in accordance with the regulations of the local ethics committee. Our volunteer cohort was sex balanced.

Human monocyte isolation and differentiation to monocyte-derived macrophages was performed as previously described^8^. Blood was drawn by venepuncture and mixed with 3.8% (w/v) trisodium citrate (Na_3_C_6_O_7_). This was layered onto Ficoll-Paque PLUS (GE Healthcare) and separated by gradient centrifugation. PBMCs were separated into PBS containing 2 mM EDTA and red blood cells were lysed using a solution of ammonium chloride (155 mM NH_4_Cl, 10 mM KHCO_3_, 0.1 M EDTA in H_2_O). Human monocytes were isolated by positive selection using CD14 microbeads (Miltenyi Biotec). Isolated monocytes were incubated in RPMI 1640 medium (Gibco) containing 10% (v/v) HI-FBS, 1% (v/v) L-Glutamine (Lonza), 1% (v/v) penicillin-streptomycin (Gibco) and 100 nM M-CSF (Peprotech) for 7-14 days at 37°C and 5% CO_2_.

### Human monocyte-derived macrophage fisetin regimen

24 hours after monocyte isolation, a 70 mM stock solution of fisetin (Stratech Scientific) suspended in DMSO (Sigma-Aldrich) was diluted (1:100) with PBS and added to the cells to give a final concentration of 10 μM. This was repeated every two days until day 14, where cells were used for functional assays and gene expression analysis, compared to vehicle control treatments using 1% (v/v) DMSO added to the cells under the same regimen.

### Phagocytosis assay

Phagocytosis was measured by uptake of Fluoresbrite YG microspheres 1.00 μm (Polysciences) and mCherry expressing *Staphylococcus aureus* strain SH1000^19^, as previously described^8^. For both mouse BMDMs and human MDMs, media was replaced with antibiotic-free media, before microspheres were opsonised in FBS for 30 minutes at 37°C and then added to each well at a concentration of 10 beads per cell. *S. aureus* was also added to the denoted wells at a multiplicity of infection of 10. Cells were then incubated at 37°C or 4°C (control) for 3 hours so that each sample had a control, microsphere and *S. aureus* well in triplicate at the two different temperatures. After 3 hours, cells were washed with PBS and fixed with 4% paraformaldehyde (Fisher Scientific) for 30 minutes at room temperature. Cells were washed again with PBS and kept in PBS at 4°C until imaging. Images were acquired on a ZOE Fluorescent cell imager (BIO-RAD) at 20x zoom, capturing a brightfield image (Gain: 5, Exposure: 340 ms, LED intensity: 59, Contrast: 20) and either a green channel (Gain: 40, Exposure: 500 ms, LED intensity: 43, Contrast: 0) or red channel (gain: 60, exposure: 500, LED intensity: 72, contrast: 0) image for each field, with three fields collected per temperature, condition and donor. Images were analysed in ImageJ v2.9.0/1.53t by applying a consistent intensity threshold to the brightfield image to generate a binary mask and using “Analyze Particles” to define regions of interest (ROIs) corresponding to individual macrophages. These ROIs were transferred onto the corresponding fluorescent image to obtain the mean gray value for each cell, and the geometric mean fluorescence per field was calculated and normalised to the matched 4°C control image before averaging across fields for each donor or mouse and condition.

### MCP-1 transwell migration assay

An MCP-1-directed transwell migration assay was performed on human monocyte-derived macrophages (hMDMs) and mouse bone marrow-derived macrophages (BMDMs) as previously described^8^. BMDMs and hMDMs were seeded at a density of 500,000 cells per mL and then collected using a cell scraper. Macrophages were centrifuged at 4000 × g for 5 minutes and stained using PKH26 red fluorescent cell linker (Sigma-Aldrich) according to the manufacturer’s instructions. Excess dye was removed by 1% BSA (Sigma-Aldrich) and cells were centrifuged at 4000 × g for 5 minutes before being resuspended at a density of 1 × 10⁶ cells/mL in 300 µL of appropriate culture media (DMEM or RPMI). Using HTS Transwell-96 permeable support (Corning), 100 µL of 1% (v/v) MCP-1 (PeproTech) in media or media alone was added to the bottom well and 100 µL of cell suspension to the transwell insert. This was incubated for 3 hours at 37 °C and 5% CO₂. After incubation, the transwell insert was removed and the bottom plate was imaged using a ZOE Fluorescent Cell Imager (Bio-Rad) at 20× magnification, with a red channel image being taken of each field and three fields being taken per sample (Gain 40, Exposure 500 ms, LED intensity 50, Contrast 20). Images were analysed in ImageJ v2.9.0/1.53t by measuring the overall mean gray value of each field. For every image, five cell-free regions were selected to obtain a mean background gray value, and background-corrected mean fluorescence values were then averaged across fields for each donor or mouse and condition.

### Bioinformatic selection of *MYC* target genes

We developed a bespoke Python (v3.10.0) based Differentially Expressed Gene Retrieval And Documentation Examination (DEGRADE) pathway, which identifies overlap between supplied RNA-sequencing datasets and chosen publicly available gene encyclopaedias, to aid in the selection of potential genes of interest for further analysis. The pipeline is publicly available on GitHub at https://github.com/lilia-elle-cs/DEGRADE.

For datasets and processing, two RNA-sequencing datasets were used from the Gene Expression Omnibus (GEO; accessed May 2023): GSE240075, containing *MYC* knockdown monocyte-derived macrophages differentiated to M₀ phenotype^8^, and GSE100905, containing CD45⁺/CD163⁺/CD11b⁺ macrophages from marrow aspirates of young and older human donors^20,21^. DEGRADE identified differentially expressed genes (DEGs) using a significance threshold of P < 0.05 with no log₂ fold-change (FC) cutoff, and classified these as upregulated (log₂FC > 0) or downregulated (log₂FC < 0).

For target gene identification DEGRADE queried the ChEA 2022 database (https://maayanlab.cloud/Harmonizome/dataset/CHEA+Transcription+Factor+Targets+2022) to extract established *MYC* transcriptional targets^22^. Intersections identified *MYC* targets concordantly dysregulated (same direction of fold-change) in both datasets (aged marrow macrophages and MYC knockdown macrophages), yielding six candidate genes with false discovery rate (FDR) < 0.05.

To select genes for RT-qPCR validation, two genes previously validated by our laboratory (GDF1 and CDH8^8^) were added to the six newly identified candidates, yielding eight genes for RT-qPCR validation (four predicted to be upregulated with age: GDF1, FCHO2, EPB41L2, DAPK1; four predicted to be downregulated with age: CDH8, SLO5A1, H1-2, YWHAG).

### RNA isolation and gene expression analysis by RT-qPCR

RNA isolation was performed using ReliaPrep RNA Cell Miniprep System (Promega) according to the manufacturer’s instructions. RNA concentration and purity was assessed using NanoDrop. iScript cDNA synthesis kit (BIO-RAD) was used to transcribe 100 ng total RNA per reaction was to cDNA in accordance with manufacturer’s instructions. RT-qPCR was performed as described previously^8^ using CFX384 C1000 Touch Thermal Cycler (BIO-RAD) and samples were assayed using Precision PLUS SYBR-Green mastermix (Primer Design). Primers were designed using NCBI BLAST tool for specific genes analysed. These sequences can be found in Supplementary Tables 1-3. All assays were performed in triplicate and normalised to expression levels of housekeeping genes PUM1 (human) or Mau2 (mouse). Relative transcript levels were analysed using 2^−ΔCt^ method.

### Quantification and statistical analysis

GraphPad Prism 10 software was used to generate all statistical analysis and graphs. N numbers and statistical analysis used are detailed for each experiment in the figure legend. Differences were considered significant where P values were < 0.05. Where possible, when comparing the same donor under different conditions, paired analysis was performed.

## Results

### Macrophage-specific Myc overexpression improves macrophage phenotype and physical performance in *Drosophila*

Previous research from our group demonstrated that *MYC* and *USF1* expression declines with age in primary human and mouse macrophages, and that MYC knockdown recapitulates age-associated functional defects including impaired chemotaxis and phagocytosis^8^. To determine whether restoring macrophage *MYC* or *USF1* expression could ameliorate functional decline *in vivo*, we utilised *Drosophila melanogaster*, a genetic model of immunity and ageing wherein plasmatocytes dominate the haemocyte blood cell lineage and function as mammalian macrophage equivalents^23–25^. *Drosophila dMyc (Myc)* is a functionally conserved orthologue of human *c-MYC (MYC)*, permitting direct *in vivo* mechanistic investigation^26,27^. *Usf* is the *Drosophila* orthologue of mammalian USF1 (https://flybase.org/reports/FBrf0218043). Consequently, we used conditional, macrophage-specific overexpression of dMyc or Usf to determine whether these evolutionarily conserved transcription factors could drive cellular changes in macrophages, similar to those observed in human cells^8^, and whether those changes in-turn would translate into improved health-span and lifespan outcomes.

Aged macrophages exhibit increased cell size compared to their younger counterparts^8^, a morphological change associated with their functional decline. To determine whether *dMyc* regulates this age-associated enlargement, we first examined *Drosophila* embryonic macrophages in which *dMyc* was specifically overexpressed (via *srpHemo-GAL4* and *crq-GAL4* drivers). Notably, *dMyc* overexpression in these cells resulted in significantly reduced whole-cell area (Figure 1A, Supplementary Figure 1A) and reduced lamellipodial area (Figure 1B, Supplementary Figure 1B) compared to controls, opposite to the increased cell size observed in aged macrophages^8^. We next assessed whether these *dMyc*-driven cellular changes translated into improved migratory function *in vivo*. Tracking embryonic macrophage migration revealed that *dMyc* overexpression significantly increased directionality during wandering migration (Figure 1C/D), showing more linear movement patterns. Enhanced directional migration with reduced cell size and altered lamellipodial morphology suggested that *dMyc* drives cellular programmes that improve macrophage function at the cellular level during development. In contrast, *Usf1* overexpression had no effect on these phenotypes (Supplementary Figure 1A-C), for this reason we focused our investigation to macrophage-specific overexpression of *Myc* and its effect on *Drosophila* lifespan.

**Figure 1.**
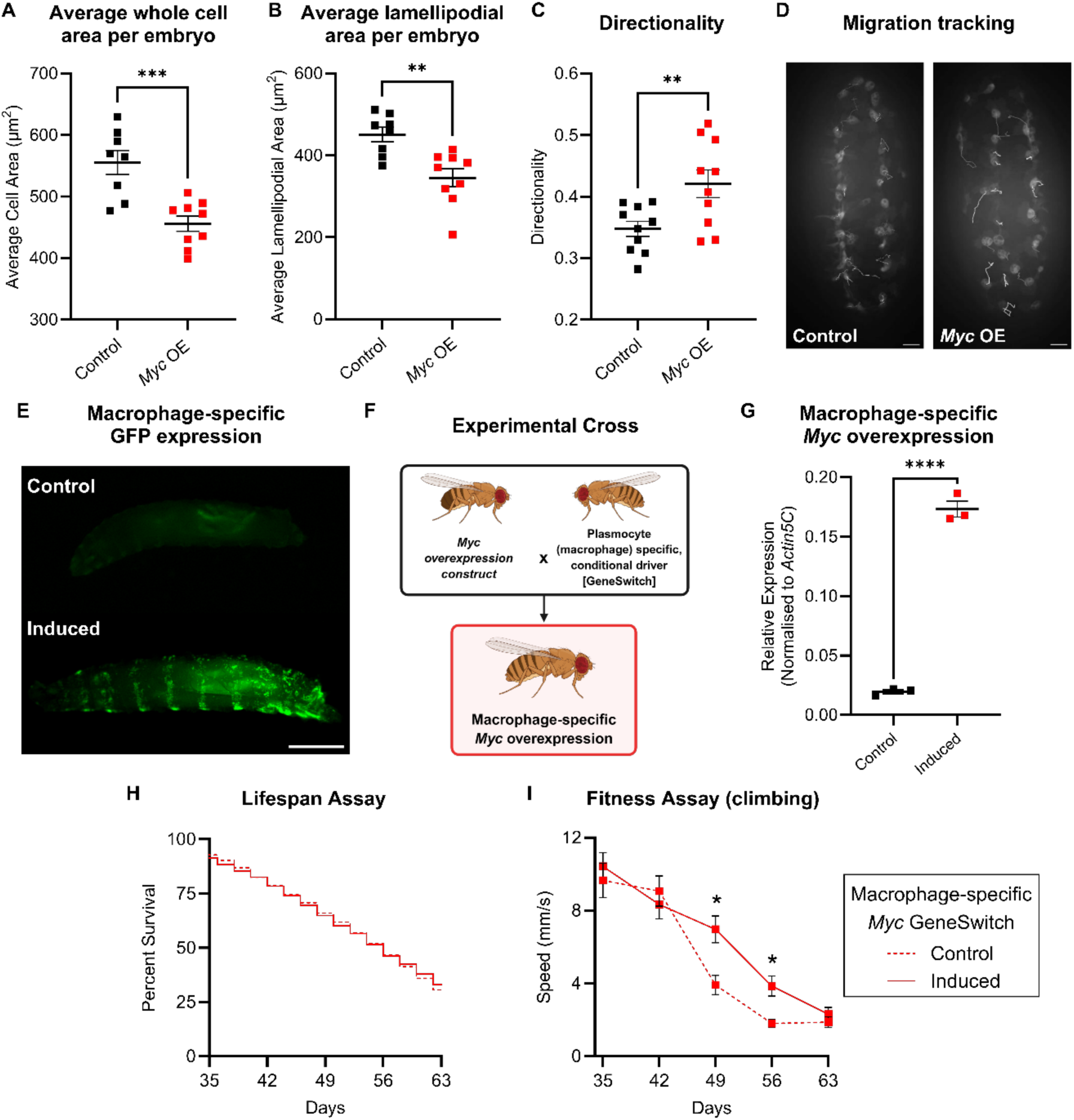
Macrophage-specific *Myc* overexpression improves macrophage phenotype and physical performance in Drosophila. **A-D** Drosophila embryonic macrophage phenotypes upon overexpression of *Myc* specifically in these cells. Live embryos containing macrophages expressing GFP +/− *Myc* expression (*Myc* OE / control) were imaged on their ventral sides at stage 15 of development. **A.** Average macrophage area per embryo (ìm²). Data are presented as mean ± SEM, with each data point representing one embryo. Unpaired t-test between control and *Myc* OE embryos, n = 8 control and n = 9 *Myc* OE, ***P < 0.001. **B.** Average lamellipodial area per macrophage, per embryo (total area minus cell body area). Data represent the mean lamellipodial area of all macrophages measured within each embryo (μm²). Data are presented as mean ± SEM, with each data point representing one embryo. Unpaired t-test between control and *Myc* OE embryos, n = 8 control and n = 9 *Myc* OE, **P < 0.01. **C.** Directionality of embryonic macrophage migration. Macrophages were tracked in vivo over 20 minutes (2-minute intervals) using the Manual Tracking plugin in Fiji. Directionality was calculated as Euclidean distance divided by accumulated distance using the Chemotaxis plugin (Ibidi) in ImageJ. Data are presented as mean ± SEM, with each data point representing one embryo. Unpaired t-test between control and *Myc* OE embryos, n = 10 per group, **P < 0.01. **D.** Migration tracking. Representative trajectories of macrophages in control embryos (left) and embryos in which *Myc* was overexpressed in macrophages (*Myc* OE, right); tracks taken from over 20-minute long movies *in vivo*. *Myc* overexpression results in more directional, streamlined migration paths. Scale bar represents 20 μm. **E.** Representative images of L3 larvae showing *Hml(Δ)-GeneSwitch* mediated expression of nuclear GFP (Stinger) specifically in the presence of RU486. Larvae fed ethanol vehicle media show minimal background fluorescence (control, upper panel), whilst those raised on RU486-supplemented media display GFP fluorescence in macrophages arranged in characteristic banding patterns along the larval body wall (induced, lower panel). Scale bar represents 1 mm. **F.** Experimental cross schematic for lifespan and climbing assay experiments. Female flies containing macrophage-specific expression of GeneSwitch (*w¹¹¹⁸;;Hml(Δ)-GeneSwitch*) were crossed with *UAS-Myc* males (*w¹¹¹⁸/Y;UAS-Myc.Z*) to generate female offspring with macrophage-specific *Myc* overexpression upon RU486 treatment (*w¹¹¹⁸;UAS-Myc.Z/*+*;Hml(Δ)-GeneSwitch/*+). Female flies were collected within a 24-hour window post-eclosion for experiments. **G.** RT-qPCR analysis of relative *Myc* expression (normalised to *Actin5C*) demonstrates macrophage-specific overexpression of *Myc* is induced via RU486. Blood cells were bled from larvae containing both *Hml(Δ)-GeneSwitch* and *UAS-Myc.Z* and maintained on vehicle control food (Control) or RU486-supplemented food (Induced). *Myc* expression was normalised to *Actin5C* (ΔCt) and plotted as fold change. Data represent pooled samples of 20 larvae from 3 independent bottles per condition. Mean ± SEM, n = 3 biological replicates, unpaired t-test, **** P < 0.0001. **H.** Lifespan assay of flies containing both *Hml(Δ)-GeneSwitch* and *UAS-Myc.Z* maintained on control media (dashed line) or RU486-supplemented media (induced, solid line;*Myc* overexpression) over 35-63 days. No significant difference in survival between groups. Logrank (Mantel-Cox) test, n = 120 flies per condition across 3 cohorts, P > 0.05. **I.** Fitness assay (climbing performance) of flies containing both *Hml(Δ)-GeneSwitch* and *UAS-Myc.Z* maintained on control media (dashed line) or RU486-supplemented media (induced, solid line; *Myc* overexpression). Median climbing speed of flies during startle-induced negative geotaxis assays at indicated time points, measured using TrackMate in Fiji. Significant improvement in climbing performance was observed at days 49 and 56. Data presented as mean ± SEM, n = 30 flies per condition at study initiation (see Supplementary Table 4 for sample size at each timepoint), mixed-effects model with Šídák’s multiple comparisons test, Adj. P < 0.05. Genotypes are as follows: *w1118;srp-GAL4,UAS-GFP*/+*;crq-GAL4,UAS-GFP/*+ (control, A-D), *w1118;srp-GAL4,UAS-GFP/UAS-Myc.Z;crq-GAL4,UAS-GFP/*+ (Myc OE, A-D), *w1118;;Hml(Δ)-GeneSwitch/UAS-stinger* (E), *w1118;UAS-Myc.Z/*+*; Hml(Δ)-GeneSwitch/*+ (G-I).

To determine whether *dMyc*-driven improvements in macrophage morphology and migratory function could extend to ageing outcomes, we utilised a conditional GeneSwitch-based system^13^ enabling macrophage-specific *Myc* overexpression throughout the fly lifecycle as controlled by the addition of the inducing drug RU486 in fly media (Figure 1F). The efficacy and specificity of the GeneSwitch system was confirmed via induction of GFP with larval blood cells in response to RU486 (Figure 1E). Furthermore, RT-qPCR analysis confirmed successful *dMyc* overexpression, with RU486-treated *Hml(Δ)-GeneSwitch/UAS-Myc* larvae exhibiting an 8.9-fold increase in *dMyc* expression compared to controls (Figure 1G).

To assess the impact of sustained macrophage *dMyc* overexpression on ageing outcomes, we measured both lifespan and physical performance of the transgenic flies. Survival analysis revealed no significant difference in lifespan between macrophage-specific *dMyc* overexpressing flies versus controls (Figure 1H; Supplementary Figure 1D), indicating that macrophage-specific *dMyc* overexpression does not extend overall survival. However, fitness assessment through climbing performance assays demonstrated a significant extension in physical performance, with d*Myc*-overexpressing flies showing enhanced climbing speed at day 49 and day 56 compared to controls (Figure 1I; Supplementary Figure 1E). These findings demonstrate that macrophage-specific *dMyc* overexpression alone improves physical performance during ageing, demonstrating for the first time a role for macrophage ageing on physical function.

### Fisetin treatment restores MYC expression and improves function in aged human monocyte-derived macrophages

Having demonstrated that macrophage-specific *Myc* overexpression improves cellular phenotype and extends health-span in *Drosophila* (Figure 1), we investigated whether the age-related decline in mammalian *MYC* expression^8^ and their corresponding functional decline could be pharmacologically reversed in human macrophages. Monocytes from young (<30 years) and older (>50 years) donors were isolated and differentiated to MDMs over 14 days in M-CSF-containing media supplemented with fisetin or equivalent DMSO vehicle control (Figure 2A). Consistent with our previous findings^8^, MYC and USF1 expression declined with age in control groups (Figure 2B). Consequently, phagocytic capacity for opsonised fluorescent beads and *Staphylococcus aureus*, as well as monocyte chemoattractant protein-1 (MCP-1) directed chemotaxis, were similarly decreased in aged controls (Figure 2C-D). Notably, fisetin selectively restored MYC expression in aged MDMs, consistent with the restoration of *MYC* (Figure 2B). In addition, fisetin-treated MDMs also showed restored phagocytic capacity and MCP-1 directed chemotactic migration (in response to fisetin treatment, Figure 2C-D). Overall, these data demonstrate that fisetin treatment restores *MYC* expression in aged primary human macrophages, with associated functional improvements, consistent with *MYC*-mediated rescue of age-related decline.

**Figure 2.**
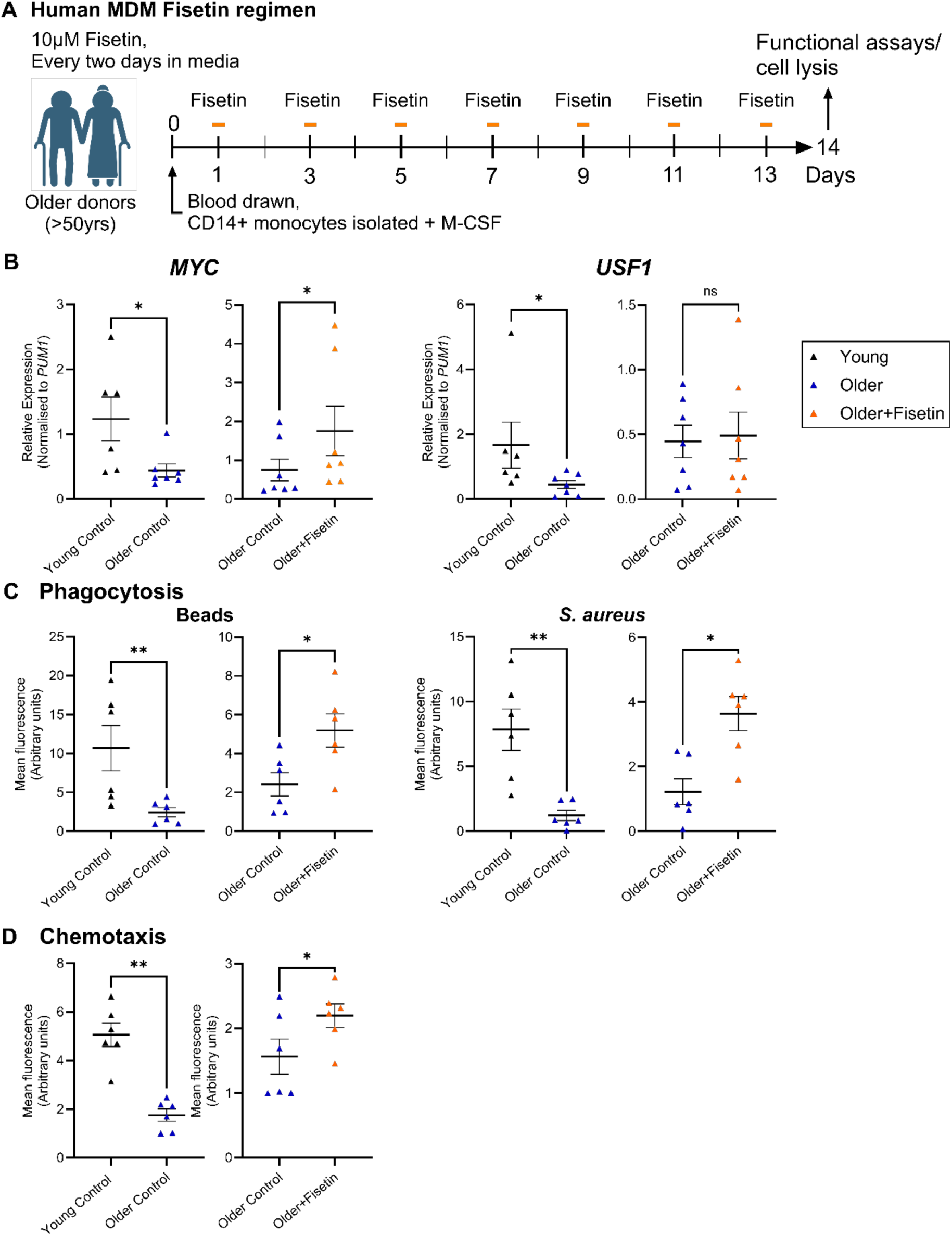
Fisetin treatment restores *MYC* expression in macrophages isolated from older humans. **A** – Schematic diagram of fisetin or vehicle control (1% (v/v) DMSO) regimen added *in vitro* to monocyte-derived macrophages isolated from older humans (>50 years). **B** – RT-qPCR analysis of *MYC* and *USF1* expression in MDMs isolated from younger (<30 years) and older (>50 years) humans and treated with fisetin *in vitro*. Older control donors in left hand graph are the same older control donors tested with fisetin. *PUM1* expression was used as a housekeeping control. Data are presented as mean ± SEM with each datapoint representing the mean of an individual donor performed in triplicate for each condition. n = 6 (young control), n = 7 (older control and older fisetin-treated), Mann-Whitney test performed between young and older controls, Wilcoxon test performed between older control and older fisetin-treated cells, * P < 0.05. **C** – Fluorescent bead uptake and *Staphylococcus aureus* uptake measured as mean fluorescence intensity (arbitrary units) after 3 hours for MDMs isolated from younger (≤30 years) and older (≥50 years) humans and treated with fisetin *in vitro*. Data are presented as mean ± SEM, with each datapoint representing the mean of an individual donor measured in triplicate for each condition, n = 6 donors. Mann–Whitney test was performed between young and older control groups, and Wilcoxon matched-pairs test between older control and older fisetin-treated cells; * P < 0.05, ** P < 0.01. MDMs were cultured for 14 days following monocyte isolation. **D** – MCP-1-directed chemotaxis measured as mean fluorescence intensity of PKH26-stained macrophages after 3 hours for MDMs isolated from younger (≤30 years) and older (≥50 years) humans and treated with fisetin *in vitro*. Data are presented as mean ± SEM, with each datapoint representing the mean of an individual donor performed in triplicate for each condition, n = 6 donors. Mann–Whitney test was performed between young and older control, and Wilcoxon test between older control and older fisetin-treated cells, * P < 0.05, ** P < 0.01. MDMs were cultured for 14 days following monocyte isolation.

### Age-related changes in expression of downstream targets of MYC are restored following *in vitro* fisetin treatment

To explore the molecular mechanisms of fisetin treatment on the *c-MYC* pathway in older macrophages, we next interrogated genes downstream of *MYC* that have an age-associated change in expression. Using our DEGRADE bioinformatic pipeline to compare aged marrow macrophages (GSE100905) and *MYC*-knockdown MDMs (GSE240075) against the ChEA database, we identified common dysregulated *MYC* targets (Table 1 and Figure 3A). Overall, six genes were found to be significantly enriched in both datasets meeting the criteria for inclusion in our experimental analysis. Our previous work^8^ identified two additional MYC target genes, affected by *MYC* knockdown in human MDMs, GDF1 and CDH8. This extended our current experimental analysis to four MYC target genes upregulated with age (GDF1, FCHO2, EPB41L2, DAPK1) and four downregulated with age (CDH8, SLO5A1, H1-2, YWHAG).

**Figure 3.**
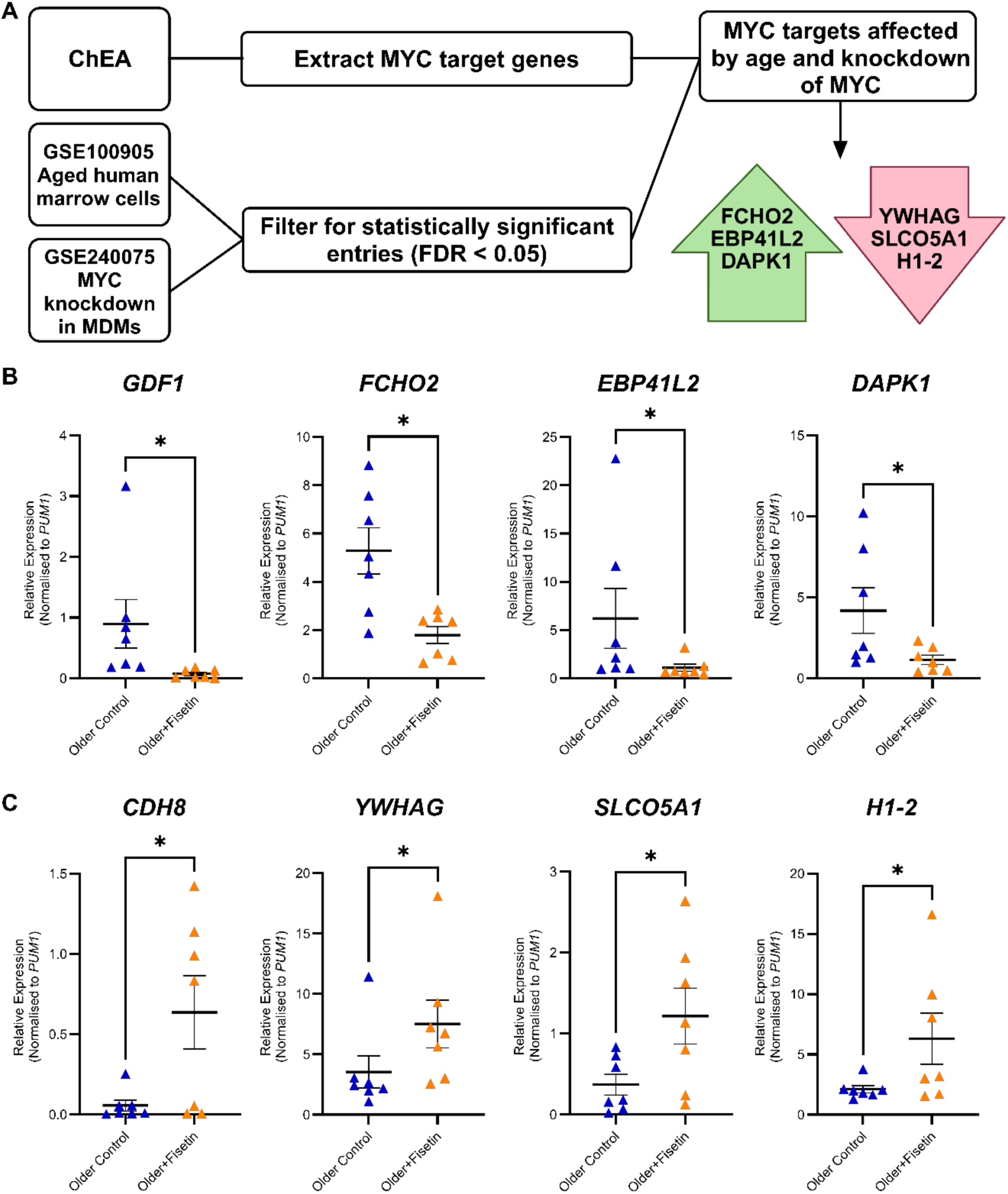
Age-related changes in expression of downstream targets of MYC are improved following *in vitro* fisetin treatment. **A** – DEGRADE pipeline to identify MYC target genes that are significantly differentially expressed in MYC-knockdown human MDMs and human marrow cells from donors of different age groups. **B,C** – RT-qPCR analysis of genes upregulated (B) or downregulated (C) with age in MDMs isolated from older (≥50 years) humans treated with vehicle control or fisetin *in vitro*. *PUM1* expression was used as a housekeeping control. Data are presented as mean ± SEM, with each datapoint representing the mean of an individual donor measured in triplicate for each condition, n = 7. Wilcoxon matched-pairs test was performed between older control and older fisetin-treated cells; * P < 0.05.

**Table 1:**
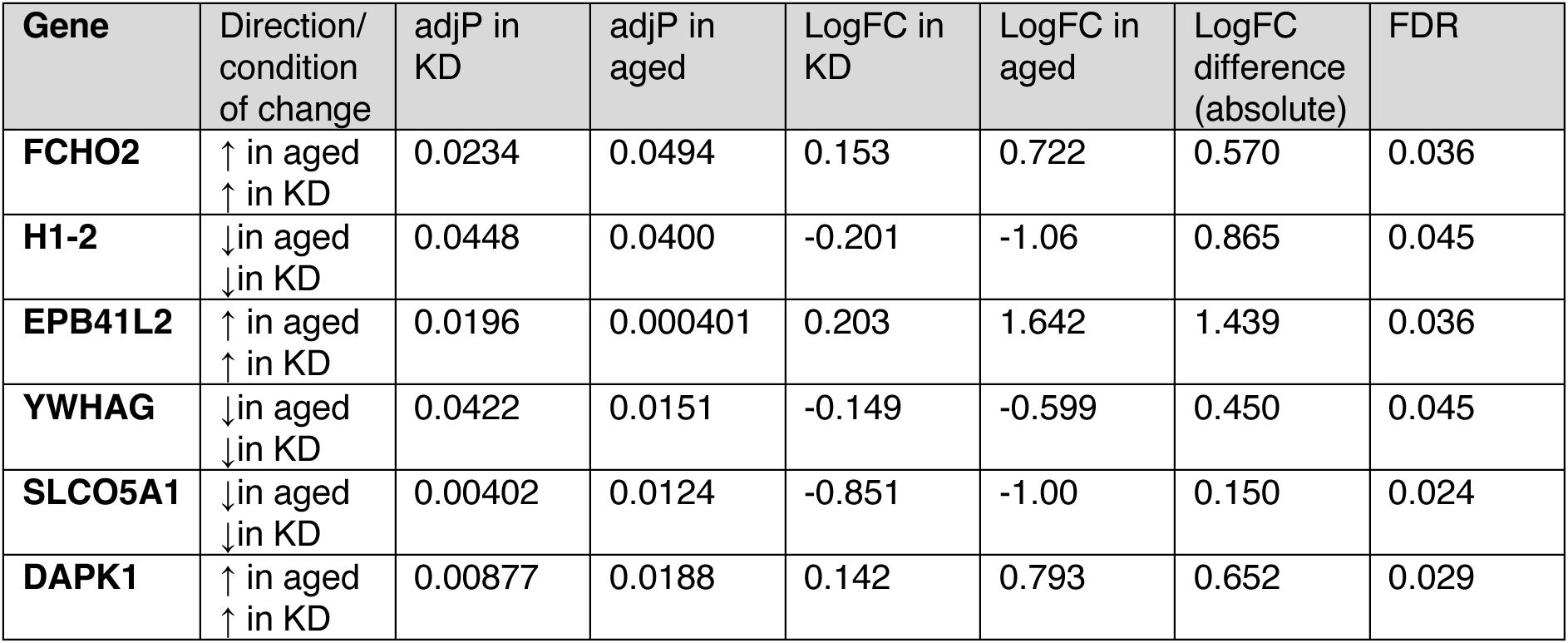
Macrophage MYC-target genes differentially expressed with age and MYC knockdown, identified using the DEGRADE analysis tool. (KD = knockdown; adjP = adjusted p-value; FC = fold change; FDR = false discovery rate)

Consistent with restoration of *MYC* levels by fisetin treatment in aged macrophages, expression of the above identified 8 targets was also significantly altered in response to fisetin, consistently restored in the same direction and to levels seen in macrophages isolated from younger donors. Thus, GDF1, FCHO2, EPB41L2, DAPK1 were significantly downregulated (Figure 3B), while H1-2, YWHAG, CDH8 and SLO5A1 were significantly upregulated following fisetin treatment (Figure 3C).

These findings demonstrate that fisetin restores the MYC-governed transcriptional program in aged macrophages, through the bidirectional reversal of dysregulation across multiple downstream targets. This functional restoration of *MYC* activity provides mechanistic insight into fisetin’s effects on macrophage phenotype.

### Fisetin remodels inflammatory and SASP marker expression in aged human macrophages

To investigate how the fisetin-induced functional restoration of MYC activity translates to macrophage phenotype, we assessed the expression of markers associated with inflammation, polarisation, and senescence-associated secretory phenotype (SASP) in unpolarised human MDMs. We compared expression in MDMs from younger and older individuals and evaluated the effect of fisetin treatment in older macrophages (Figure 4).

**Figure 4.**
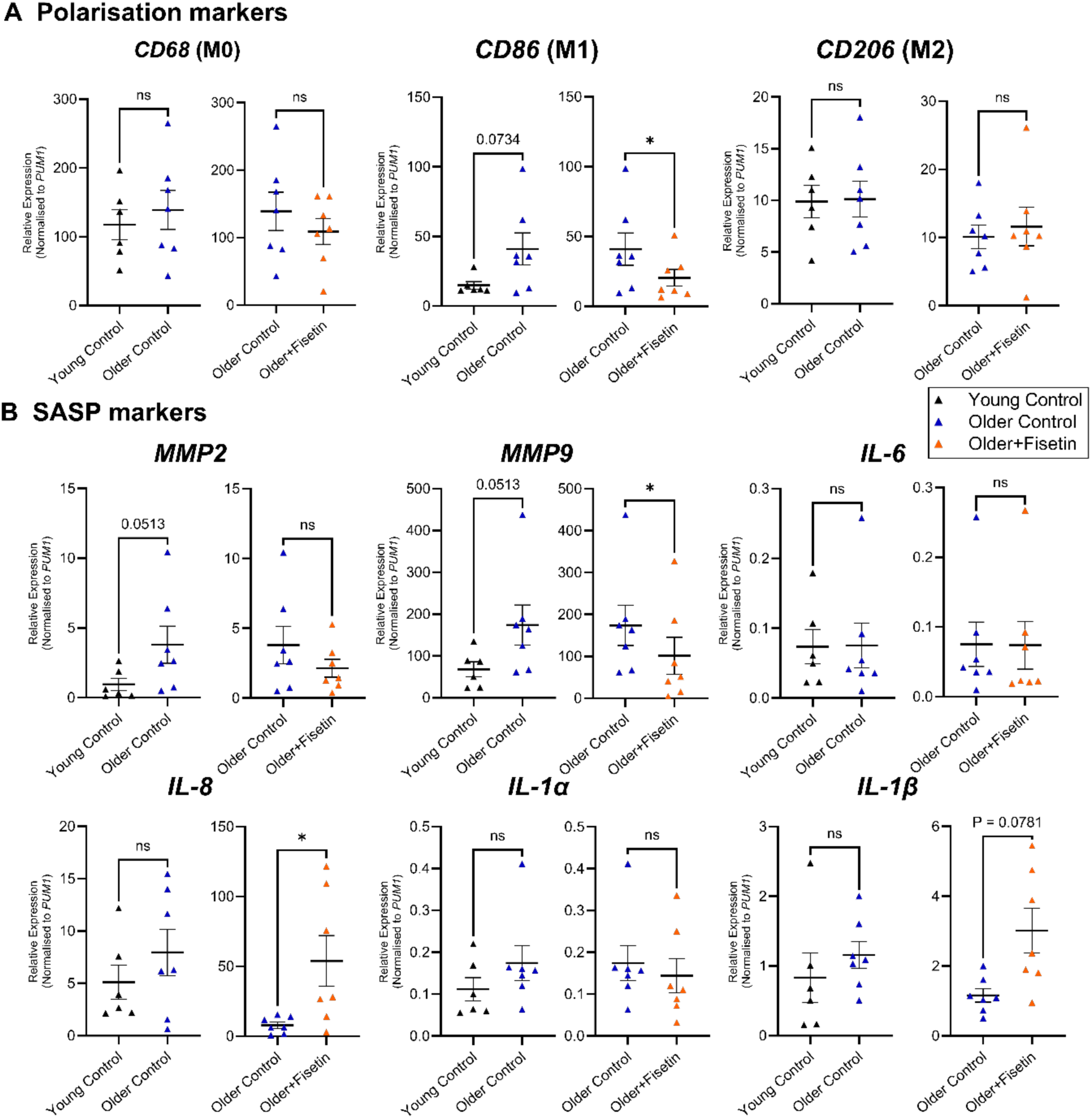
Effect of Fisetin on Polarisation and SASP markers in macrophages from older humans. RT-qPCR analysis of polarisation markers (**A**) and SASP markers (**B**) in MDMs isolated from young (<30 years) and older (>50 years) humans and treated with fisetin *in vitro*. *PUM1* expression was used as a housekeeping control. Data are presented as mean ± SEM with each datapoint representing the mean of an individual donor performed in triplicate for each condition, n = 6-7. Mann-Whitney test performed between young and older control, Wilcoxon test performed between older control and older fisetin-treated cells, included in one graph for better visualisation, * P < 0.05.

Among polarisation markers, CD86, a marker of M1 pro-inflammatory polarisation, displayed an upward trend with age (p = 0.0734) and was significantly downregulated by fisetin treatment in older macrophages (Figure 4A). Examination of SASP markers revealed selective remodelling of the inflammatory secretome (Figure 4B). MMP9 expression was significantly reduced by fisetin treatment in older macrophages (Figure 4B), whilst IL-8 expression was significantly increased by fisetin treatment (Figure 4B), with IL-6 and IL-1β expression not altered by age or fisetin. The selective downregulation of CD86 and MMP9 with increased IL-8 suggests a complex remodelling of the macrophage inflammatory phenotype that extends beyond conventional M1/M2 polarisation.

### Fisetin improves macrophage function and health-span in aged, multimorbid mice

Having established that fisetin selectively restores *MYC* expression and function in aged human MDMs *in vitro* (Figures 2-3), we next investigated whether these cellular changes translate to improved health-span in aged atherosclerotic mice. These mice were aged to 16 months and then administered a Western diet containing fisetin over a 12-week period (Figure 5A). Bone marrow was isolated and cultured in M-CSF for 5 days, with no further fisetin supplementation *in vitro,* to generate bone marrow-derived macrophages (BMDMs), which were either left unstimulated (M^0^), polarised towards a pro-inflammatory phenotype with lipopolysaccharide (LPS) and interferon-γ (M^LPS+IFNγ^), or polarised towards an anti-inflammatory phenotype with interleukin-4 (M^IL-4^). In unpolarised M0 macrophages, where *Myc* expression was previously shown to decline with age^8^, fisetin treatment restored *Myc* expression compared with aged controls (Figure 5B). The restoration of *Myc* expression led to improved macrophage function, where BMDMs from fisetin supplemented mice exhibited significantly improved *Staphylococcus aureus* uptake (Figure 5C).

**Figure 5.**
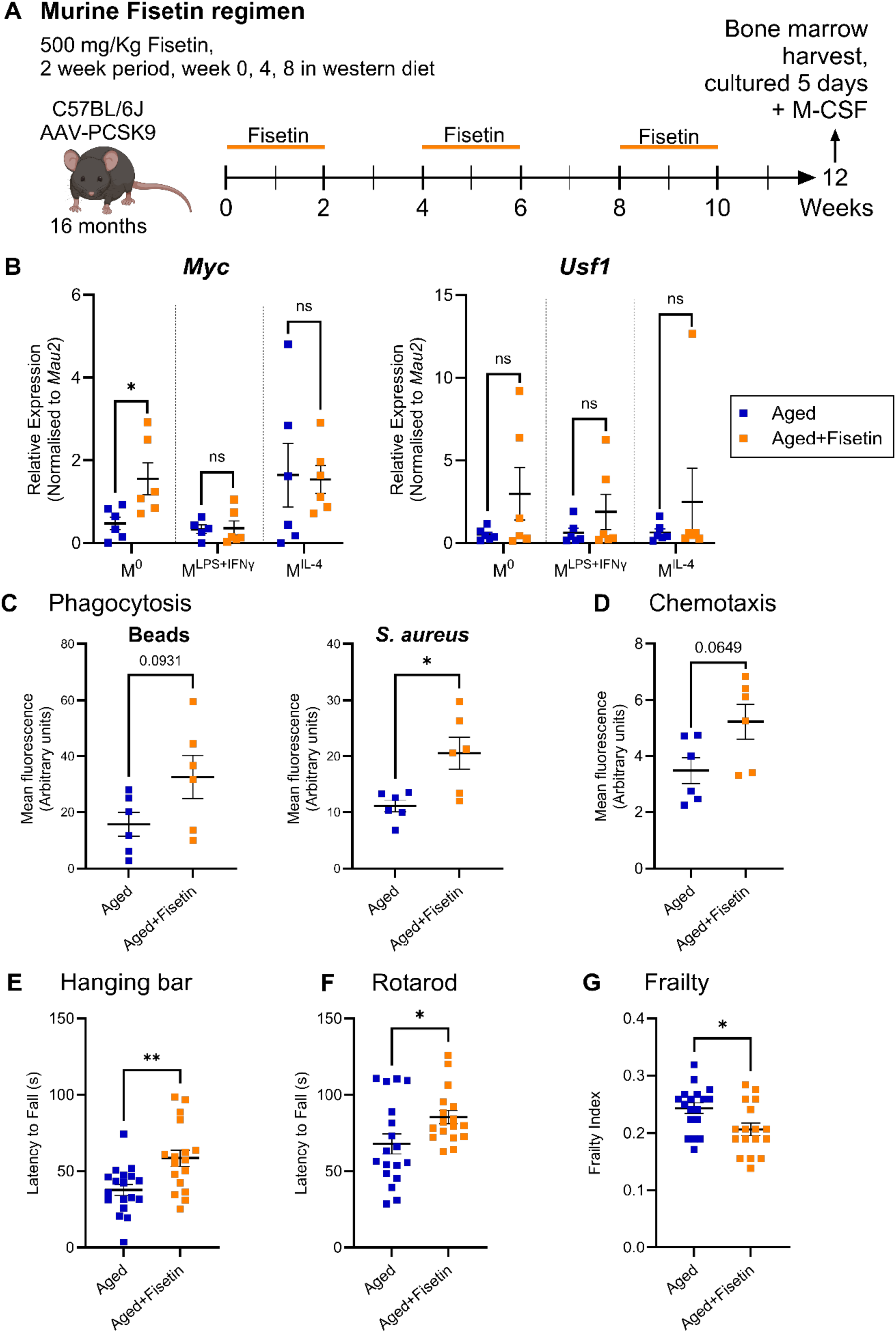
Fisetin improves macrophage function and health-span in aged atherosclerotic mice. **A** – Schematic diagram of the fisetin supplemented (500 mg/kg/day) western diet (WD) given orally to rAAV/D377Y-mPCSK9-C57BL6/J mice aged 16 months for 2 weeks every month for 3 months. Control aged (20 months) mice were fed a WD only. **B** – RT-qPCR analysis of *Myc* and *Usf1* expression in BMDMs isolated from aged mice that were fed a WD and aged mice fed a WD supplemented with fisetin. BMDMs were isolated and cultured for 5 days following bone marrow extraction. M^0^, cells unstimulated; M^LPS+IFNγ^, cells stimulated with100 ng/ml LPS and 20 ng/ml IFN-γ for 24 h; M^IL-4^, cells stimulated with 20ng/ml IL-4 for 24 h. *Mau2* expression was used as a housekeeping control. Data are presented as mean ± SEM, with each datapoint representing the mean of an individual mouse performed in triplicate for each condition, n = 6, Mann–Whitney test, * P < 0.05. **C** – Fluorescent bead uptake and *Staphylococcus aureus* uptake measured as mean fluorescent intensity after 3 hours for BMDMs isolated from aged mice and aged mice fed a WD supplemented with fisetin. Data are presented as mean ± SEM, with each datapoint representing the mean of triplicate experiments done per mouse, n = 6, Mann–Whitney test, * P < 0.05. **D** – MCP-1-directed chemotaxis measured as mean fluorescence intensity of PKH26-stained macrophages after 3 hours for BMDMs isolated from aged mice and aged mice supplemented with fisetin. Data are presented as mean ± SEM, with each datapoint representing the mean of an individual mouse performed in triplicate for each condition, n = 6, Mann–Whitney test, * P < 0.05. **E** – Grid hanging bar test performance measuring muscle strength and endurance. Data are presented as mean ± SEM, with each datapoint representing an individual mouse, n = 18 aged and n = 17 aged + fisetin, unpaired two-tailed t-test, ** P < 0.01. **F** – Rotarod performance assessing motor coordination and balance. Data are presented as mean ± SEM with each datapoint representing an individual mouse. n = 18 (aged) and n = 17 (aged + fisetin), unpaired two-tailed t-test, * P < 0.05. **G** – Frailty index calculated based on 29 standardized health deficits. Data are presented as mean ± SEM with each datapoint representing an individual mouse. n = 18 (aged) and n = 17 (aged + fisetin), unpaired two-tailed t-test, * P < 0.05.

To determine if these cellular improvements by fisetin translate to health outcomes, we assessed physical performance and frailty in aged mice at 20 months. Fisetin-supplemented aged mice showed improved muscle strength and endurance on the grid hanging bar test (Figure 5E), improved motor coordination and balance on rotarod (Figure 5F), and significantly reduced frailty index scores (Figure 5G), demonstrating that fisetin-mediated restoration of MYC-dependent macrophage function improves physical performance and reduces age-related frailty.

## Discussion

In this study we show that fisetin-mediated restoration of MYC expression in macrophages from older individuals improves immune function and health-related motor activity. Macrophage-specific *Myc* overexpression in *Drosophila* and fisetin-induced upregulation of Myc improves physical performance in older individuals, while lifespan was unaffected. In primary human macrophages from older individuals, fisetin selectively restored *MYC* and *MYC* target gene expression and recovered age-associated functional decline. Collectively, these results demonstrate that restoring MYC expression through either genetic overexpression or pharmacological intervention reverses age-associated macrophage dysfunction across multiple model systems and improves physical performance.

Macrophages play a key role in inflammaging and age-associated immune decline. Most work to date on macrophage ageing has been focussing on analysis of mouse tissues. Recently we found that human macrophages show a profound reduction in phagocytosis, migration and chemotaxis in the older age group, driven by key transcriptional regulators MYC and USF1^8^. We therefore assessed the effect of macrophage-specific USF1 and MYC overexpression on macrophage phenotype, lifespan and physical performance, an important index of health, using an *in vivo Drosophila* model. While *Usf1* overexpression did not appear to alter these measures, macrophage-specific *dMyc* overexpression resulted in a significant reduction in cell size, reduced lamellopodial area and an improvement in migration directionality. This aligns to our previous findings where loss of *MYC* in human monocyte-derived macrophages either with age or by knockdown, results in reduced migration, increased cell size and reduced circularity^8^. The *Drosophila* model provides an effective tool to assess, for the first time, the effect of macrophage (plasmocyte)-specific *Myc* overexpression on lifespan. Here we found that overexpression of *Myc* in these cells did not significantly affect lifespan in each of the control and *Myc* overexpressing cohorts; there was no obvious tumour development in this experimental cohort. This lack of effect on lifespan contrasts with studies showing that reduced expression of *Myc* in heterozygous mouse knockouts increases longevity and enhances lifespan^28^; it is important to note however, that these mouse studies used a whole-body knockout approach. Whole-body overexpression of *Myc* in Drosophila has been shown to cause genome instability and a reduction in lifespan^29^. Our work is mechanistically more refined, since *Myc* overexpression was targeted specifically to macrophages, which then did not adversely affect lifespan.

Macrophage-specific *Myc* overexpression in *Drosophila* improved their fitness assessed by climbing performance, but only in the older-aged individuals, between days 49 to 56 which roughly associates with human age of around 80 years^30^. Our human and mouse studies have shown that macrophage levels of MYC decline with age, with the fly model suggesting that restoring expression in macrophage alone can improve physiological function and fitness. The improved fitness in flies fits with our additional findings here, where older mice show improved hanging bar and rotarod fitness when macrophage *Myc* expression is upregulated as a result of dietary fisetin supplementation.

Fisetin, a naturally-occurring flavonoid found in several fruits, has emerged as a promising geroprotective compound^16^. Its administration to wild-type mice late in life restores tissue homeostasis, reduces age-related pathology, and extends median and maximum lifespan^16^. Preclinical studies have demonstrated fisetin’s protective effects across multiple cell types and tissues relevant to ageing. In aged mice, intermittent fisetin supplementation improves arterial function by decreasing cellular senescence and reducing vascular inflammation^31^.

In addition to its senolytic properties, fisetin possesses anti-inflammatory, antioxidant, anti-cancer and anti-hyperglycaemic activities^16,32,33^. Fisetin reduces tumour cell viability^34^, protects against c-MYC phosphorylation^35^ and is also a candidate ligand for binding DNA-G4 quadruplexes^36^, found in the c-myc promoter^37^. Notably, fisetin is currently undergoing Phase 2 clinical trials for reducing inflammation and frailty markers in ≥ 70-year-olds (AFFIRM trial, NCT03430037), and in breast cancer survivor health in post-menopausal women (TROFFi trial, NCT05595499). There remains a gap in our understanding specifically of the effect of fisetin on age-related decline in macrophage function.

We assessed whether fisetin was effective in altering key regulators of macrophage function that decline with age, since this could offer a therapeutically relevant opportunity to treat age-related disease including immune decline. We found that *MYC* was upregulated in human monocyte-derived macrophages from older individuals following *in vitro* treatment with fisetin. Macrophage function was markedly improved by fisetin treatment with respect to phagocytosis of beads and bacteria and to migration. Fisetin had no effect on improving these functions or on upregulating *MYC* expression in macrophages from younger individuals. This parallels with the finding that *MYC* upregulation in *Drosophila* macrophages showed functional improvement with respect to climbing fitness only in older individuals. The fact that fisetin does not induce upregulation of *MYC* in macrophages from younger individuals and restores levels towards that of young in older-derived macrophages, suggests its action is to restore levels rather than activate above a healthy physiological state.

To date the effect of fisetin on macrophages has been demonstrated only in cell line models. This is the first report on the effect of fisetin in primary human macrophages and comparing its effect on age-associated declining macrophage function. Fisetin was previously shown to reduce LPS-mediated inflammation in mouse peritoneal macrophages^38^, IFNγ-mediated inflammation in RAW264.7 cell line macrophages^39^ and to attenuate recruitment of immune cells following LPS injection into zebrafish larvae^40^. Here we compared the effect of fisetin on the production of inflammatory and senescence associated secretory factors in primary human macrophages from younger and older individuals. A reduction in the inflammatory macrophage marker CD86 and in MMP9 was observed in older-derived macrophages, but not from young-derived macrophages, while pro-inflammatory IL-8 was upregulated by fisetin. This suggests that fisetin is not acting simply at the conventional M1/M2 axis or in one direction regarding the SASP secretome, which may reflect its action on the MYC axis, and downstream MYC-target genes.

The transcription factor MYC is a key regulator of macrophage function that is dysregulated with age, where previously we used RNA-seq to assess changes in gene expression as a result of *MYC* knockdown in human MDMs from younger individuals^8^. To gain mechanistic insight in the current study, we employed a bioinformatic pipeline to identify which of the ChEA database MYC targets are both differentially expressed genes with age in a human macrophage dataset and by *MYC* knockdown in young hMDMs. From this we identified four *MYC* target genes upregulated with age (*GDF1, FCHO2, EPB41L2, DAPK1*) and four downregulated with age (*CDH8, SLO5A1, H1-2, YWHAG*). In human macrophages from older individuals, fisetin significantly altered expression of each of these genes in the direction of expression seen in macrophages from the younger group. Interestingly, genetic and cellular analyses in humans strongly associate genetic variants of these genes with the specific molecular pathologies that underpin frailty and the development of multiple chronic conditions.

DAPK1 is a calcium/calmodulin-regulated serine-threonine kinase involved in autophagy, endocytosis, vesicle transport, nutrient uptake and homeostasis^41^. DAPK1 activates NF-κB and NLRP3-mediated inflammation in human chondrocytes^42^. Human variants of DAPK1 are associated with altered vaccine responsiveness^43^, increased risk of cardiovascular disease and type-2 diabetes^44^. This, alongside our findings that macrophage *DAPK1* is a downstream *MYC* target that is upregulated with age suggests its dysregulation may be coupled to inflammaging. GDF1 activates Akt^45^, a kinase regulating macrophage polarisation^46^ and component of the PI3K/AKT/mTOR pathway which plays a key role in biological ageing including SASP production^47^. Fisetin treatment was effective in reducing *DAPK1* and *GDF1* expression in MDMs from older individuals suggesting its action at the MYC axis may be effective in suppressing pathways linked to age-related disease and dysfunction.

Linker histone H1-2 is required to maintain nucleosome and chromatin stability and was recently identified as a negative regulator of cGAS^48^, the cytoplasmic DNA sensor that activates the STING IFN-I inflammatory pathway. Our finding that macrophage *H1-2* is a MYC target whose levels decrease with age suggests this could be linked to chromatin instability and subsequent overactivation of the DNA sensing STING pathway that occurs in cell senescence. We found that fisetin treatment of hMDMs from older individuals upregulates *H1-2* expression, suggesting a new MYC-targeted mechanism of action for fisetin that reduces senescence. The Framingham study showed a *CDH8* variant rs2639889 is associated with morbidity-free survival at age 65^49^, although there is limited knowledge on the role of this cadherin in immunity and ageing.

YWHAG is proposed to play a role in mitophagy^50^. Linked to our findings that macrophage *YWHAG* is a MYC target showing decreased expression with age, reduced levels of *YWHAG* were recently shown to link to inflammatory macrophages and to be predictive of neurodegenerative age-associated Alzheimer’s Disease^50^. SLCO5A1 is a solute carrier transport protein, proposed to play a role in transport of chemokines and metabolites involved in the reorganisation of cell shape^51^. Fitting with our finding that macrophage *SLCO5A1* is a MYC-target gene that declines with age, downregulation of SLCO5A1 protein levels were identified in mild osteoarthritis human knee joints compared to healthy joints, suggesting this transport protein is important for a healthy joint and reduced levels are associated with inflammatory joint damage^52^. Fisetin was effective in upregulating each of these MYC-targets that decline with age and may be implicated in inflammaging-associated diseases.

Here we show that age-related function is restored with macrophage-specific Myc overexpression *in vivo* using *Drosophila* and in human primary monocyte-derived macrophages treated with fisetin acting at the *MYC* axis. Taking these results together, we assessed the effect of fisetin supplementation by feeding in an aged mouse atherosclerosis model. This model reflects a therapeutically relevant setting with respect to administration and to improving prevalent cardiovascular disease where age and frailty are major risk factors^53,54^. Fisetin significantly reduced frailty index in mice, compared to control. In primary bone-derived macrophages from mice fed fisetin, *Myc* expression was upregulated and bacterial phagocytosis was significantly increased. Similar to *Drosophila* macrophage-*Myc* overexpression, fisetin improved fitness as assessed for muscle strength and endurance by grid hanging bar and for coordination and balance by accelerating rotarod. This indicates that macrophage MYC upregulation improves fitness associated with health-span across species and identifies a new mechanism for the action of fisetin in improving biological ageing.

In summary, we have highlighted that restoring *MYC* expression either by genetic overexpression or pharmacological intervention reverses age-associated macrophage dysfunction in human primary cells, across multiple species ageing models and improves physical performance and health-span outcomes. Fisetin restores the key regulatory factor MYC that declines with age in macrophages, as well as its downstream regulated genes. Restoring the macrophage MYC axis represents a new mechanism for the action of fisetin in reducing frailty and immune ageing.

### Limitations of the study

Our studies using human MDMs were from two gender-balanced healthy volunteer age groups: 18–30 years (mean age 22.5 ± 1.8 SD years; young cohort, 7 male and 5 female) or >50 years (mean age 62 ± 6.3 SD years; older cohort, 7 female and 3 male). The healthy ageing donors are without comorbidities and frailty seen in the ageing human population, which will form more complex and interesting future studies beyond the scope of this work. Human macrophage function was determined in vitro, since live phagocytosis and migration detection would not have been possible in vivo. Assessing function and gene expression under resting and specific stimulation conditions offers a simplified model compared to macrophages in vivo. Mouse macrophages and fitness tests were from male animals, but these were extrapolated to human macrophage assessment from gender-balanced groups.

## Acknowledgements

We thank Carl Wright, Laura Martinez-Campesino and Taewoo Kim for assistance with bone marrow collections. We are grateful to Simon Johnston for supplying fluorescent-labelled *S.aureus*. We also thank phlebotomists Erin Card, Mazen Alhommrani and Sangam Gurung as well as the volunteers who donated blood to this research. We thank Kath Whitley and technicians of the University of Sheffield Fly Facility for molten food production. *Drosophila* work would not be possible without access to Flybase; stocks obtained from the Bloomington *Drosophila* Stock Center (NIH P40OD018537) were used in this study. This work was supported by the Healthy Lifespan Institute, University of Sheffield, the Vivensa Foundation and the British Heart Foundation (FS/PhD/25/29714).

## Author Contributions

JVK, CEM, LSH, MLC, IRE and AR performed experiments and data analysis. HLW and EK-T designed and supervised the study and contributed to data analysis and interpretation. IRE, RJHW, SEF and IB designed and supervised the study. JVK and HLW wrote the manuscript. All authors reviewed and edited the manuscript.

---

**Supplementary Table 1.**
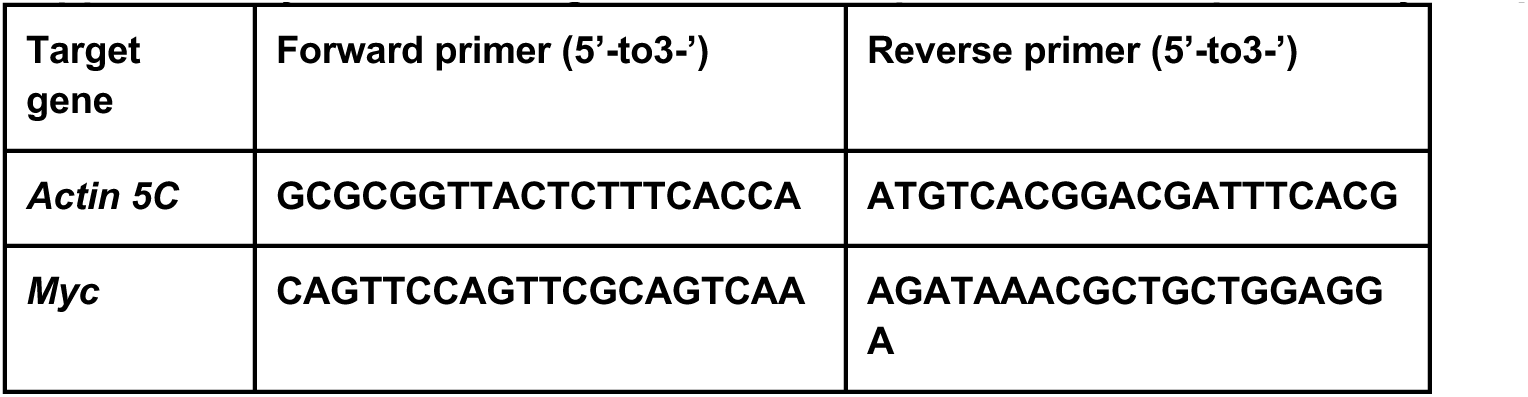
Oligonucleotide sequences for RT-qPCR of fly samples.

**Supplementary Table 2.**
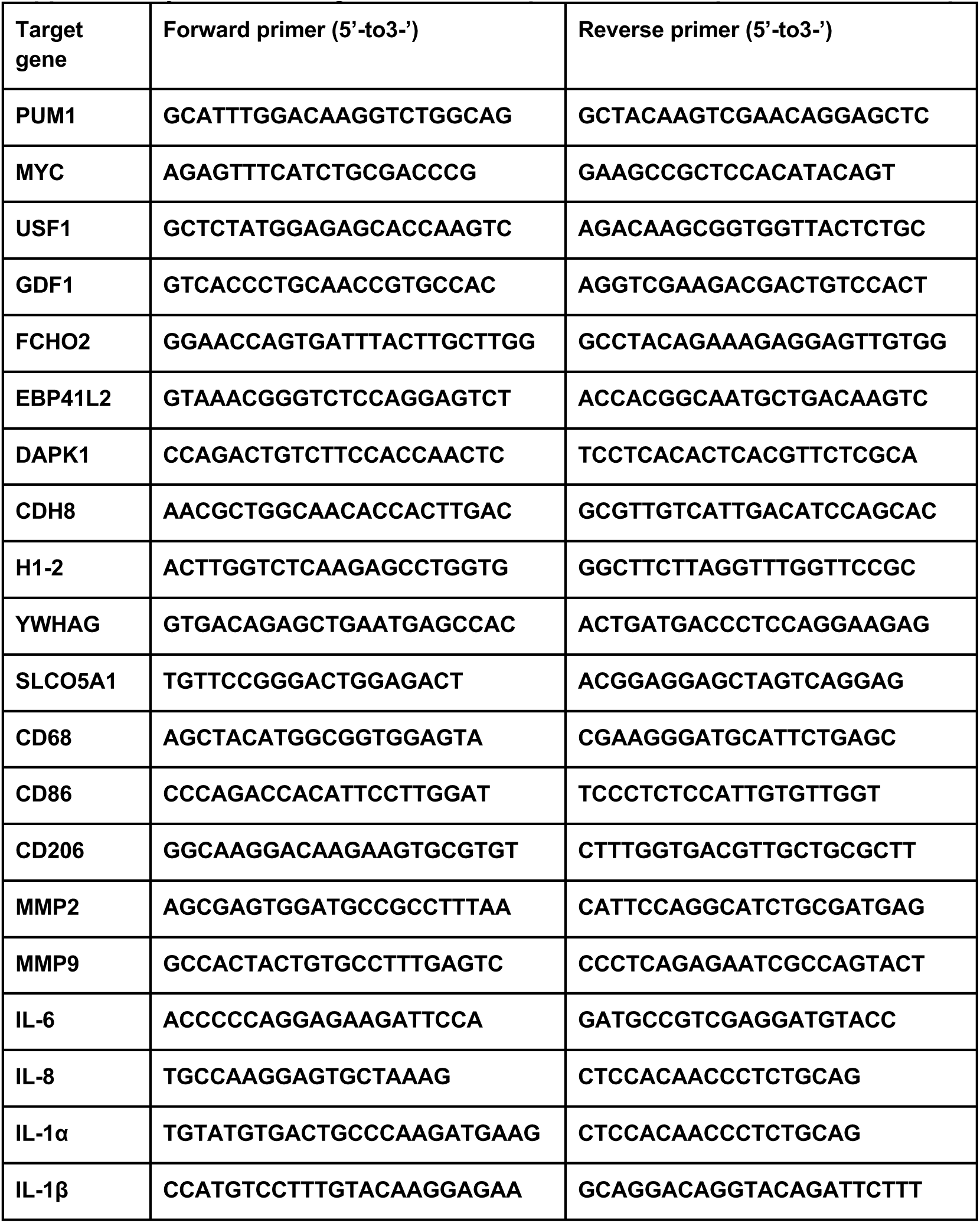
Oligonucleotide sequences for RT-qPCR of human samples.

**Supplementary Table 3.**
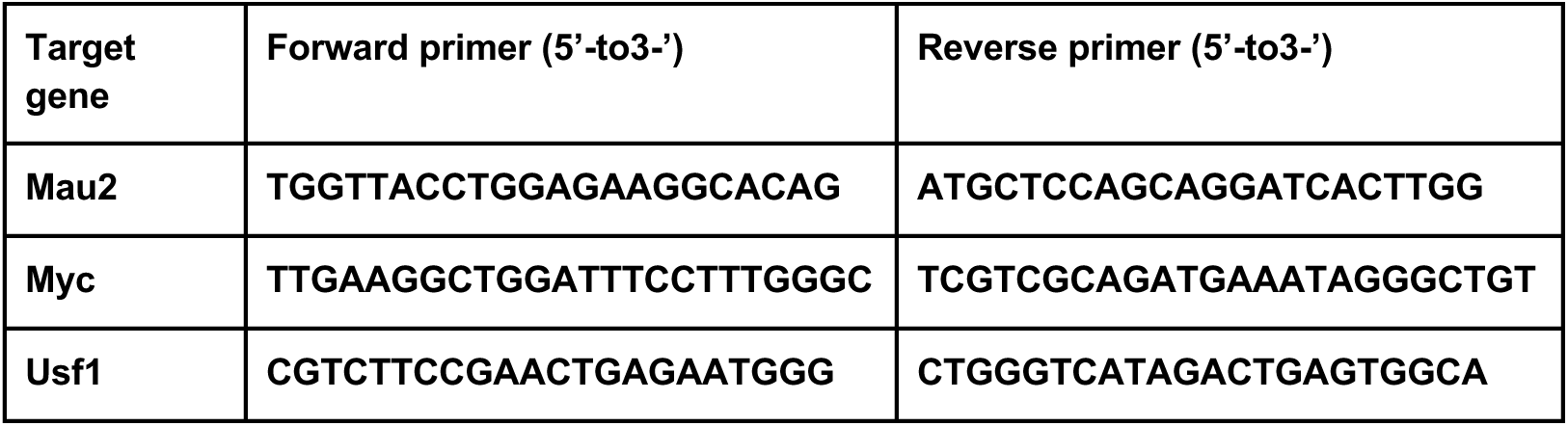
Oligonucleotide sequences for RT-qPCR of mouse samples.

**Supplementary Table 4.** Composition of standard *Drosophila* culture medium. All ingredients were mixed and autoclaved. Medium was cooled to 50-55°C before vehicle/drug added and subsequent dispensing into vials or bottles.

| Ingredient | Amount | Manufacturer | Supplier |
| --- | --- | --- | --- |
| Cold tap water | 1 Litre | - | - |
| Medium Cornmeal | 80 g | Triple Lion | Lembas/Easton Enterprises, Sheffield, UK |
| Dried Yeast | 18 g | Kerry Ingredients | BTP Drewitt, London, UK |
| Soya Flour | 10 g | Lembas Wholefoods | Lembas |
| Malt Extract | 80 g | Rayner's Essentials | Lembas |
| Molasses | 40 g | Rayner's Essentials | Lembas |
| Agar | 8 g |  | BTP Drewitt |
| 10% Nipagin in | 25 ml | Clariant UK Ltd | Chemlink Specialities Ltd, Manchester, UK |
| Absolute Ethanol |  | Fisher | Fisher |
| Propionic Acid | 4 ml | Fisher | Fisher |

**Supplementary Table 5.**
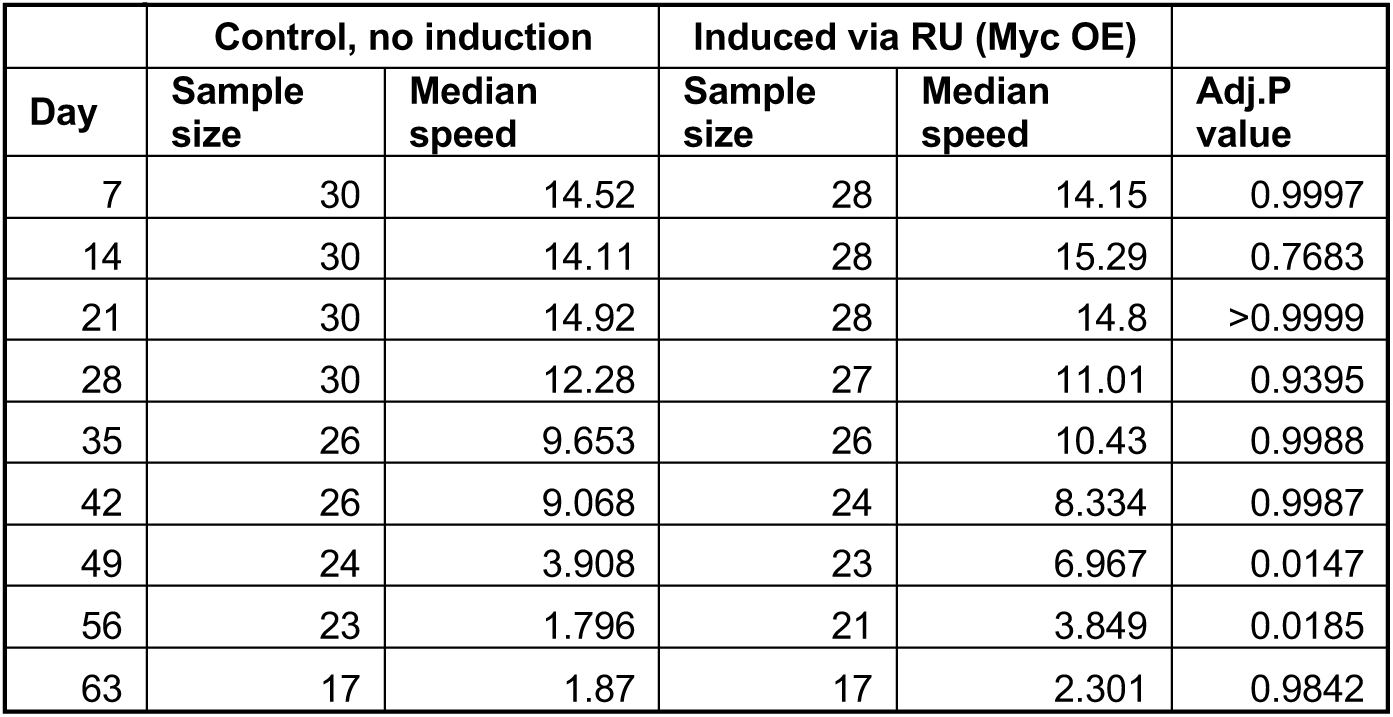
Sample sizes for climbing (negative geotaxis) assays across timepoints in *Myc* O/E control and *Myc* O/E RU486 cohorts. *n* = 30 flies per condition at study initiation, Mixed-effects model with Šídák’s multiple comparisons test, Adj.P < 0.05.

|  | Control, no induction |  | Induced via RU (*Myc* OE) |  |  |
| --- | --- | --- | --- | --- | --- |
| Day | Sample size | Median speed | Sample size | Median speed | Adj.P value |
| 7 | 30 | 14.52 | 28 | 14.15 | 0.9997 |
| 14 | 30 | 14.11 | 28 | 15.29 | 0.7683 |
| 21 | 30 | 14.92 | 28 | 14.8 | >0.9999 |
| 28 | 30 | 12.28 | 27 | 11.01 | 0.9395 |
| 35 | 26 | 9.653 | 26 | 10.43 | 0.9988 |
| 42 | 26 | 9.068 | 24 | 8.334 | 0.9987 |
| 49 | 24 | 3.908 | 23 | 6.967 | 0.0147 |
| 56 | 23 | 1.796 | 21 | 3.849 | 0.0185 |
| 63 | 17 | 1.87 | 17 | 2.301 | 0.9842 |

**Supplementary Figure 1.**
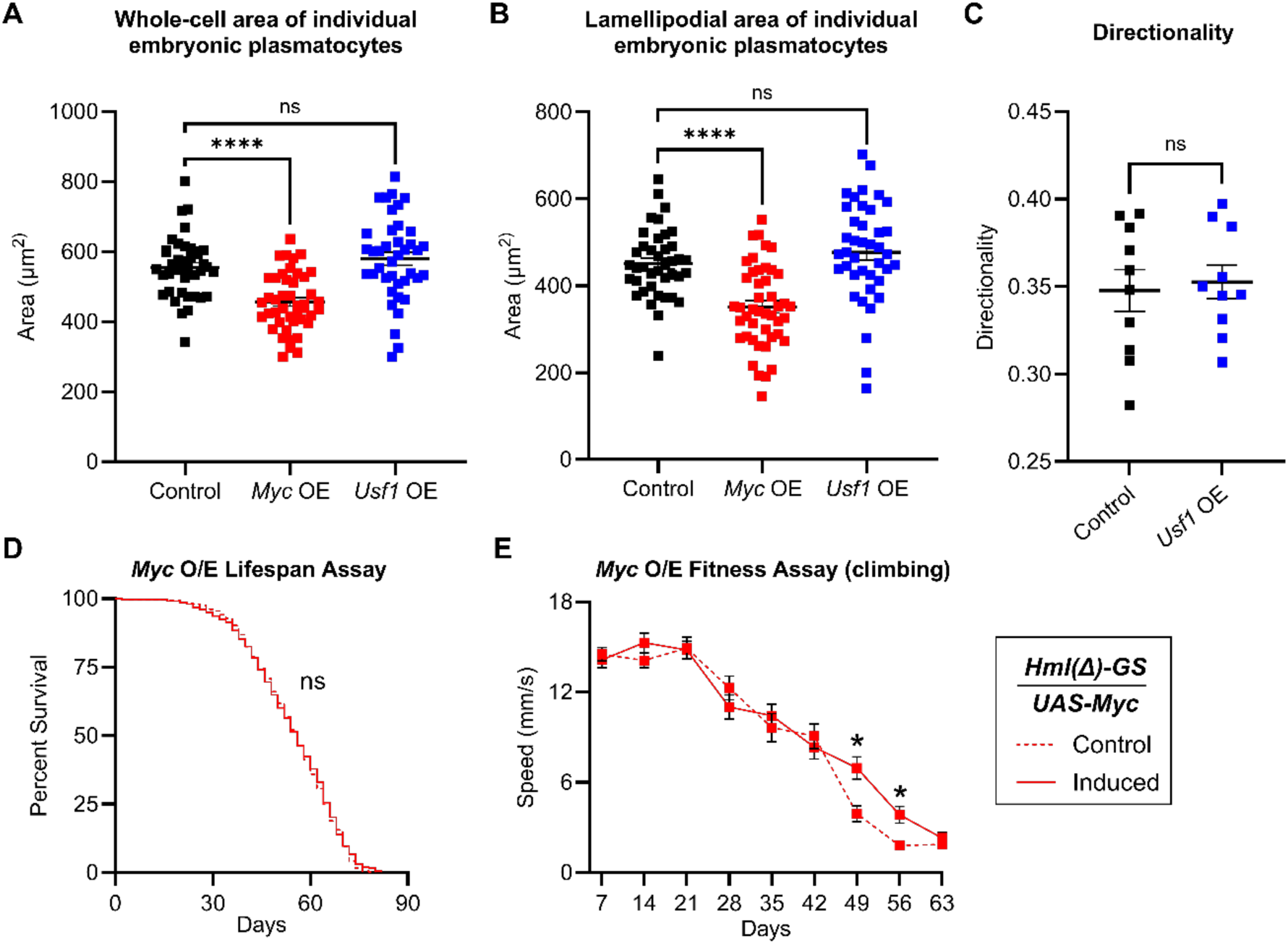
Myc-dependent modulation of *Drosophila* macrophage morphology and ageing phenotype. **A**. Individual whole-macrophage area measurements in control embryos and on macrophage-specific overexpression of *dMyc* (*Myc* OE) or *Usf* (*Usf* OE). Maximum intensity projections were generated from 30 μm z-stacks and whole-cell area was quantified in Fiji All individual cells are plotted (each point = one macrophage); data are presented as mean ± SEM, n = 40 control, n = 44 Myc OE and n = 40 Usf OE cells. One-way ANOVA with Dunnett’s multiple comparisons test was conducted between control vs *Myc* OE and *Usf* OE. **B**. Individual lamellipodial area measurements for embryonic macrophages. Lamellipodial area was calculated as whole-cell area minus cell-body area for each macrophage using the same imaging parameters and embryos described for whole-cell area analysis. All individual cells are plotted (each point = one macrophage); data are presented as mean ± SEM, n = 40 control cells, n = 45 Myc OE and n = 40 Usf OE cells. One-way ANOVA with Dunnett’s multiple comparisons test was conducted between control vs *Myc* OE and *Usf* OE. **C**. Average migration directionality per macrophage, per embryo in control and *Usf* OE embryos. Macrophages were tracked *in vivo* over 20 minutes (2-minute intervals) using the Manual Tracking plugin in Fiji. Directionality was calculated as Euclidean distance divided by accumulated distance using the Chemotaxis plugin (Ibidi) in ImageJ. Data are presented as mean ± SEM, with each data point representing one embryo, n = 10 embryos. Unpaired t-test between control vs. *Usf* OE (*n* = 10) embryos, P > 0.05. **D** Survival curves of flies containing both *Hml(Δ)-GeneSwitch* and *UAS-Myc.Z* maintained on control media (dashed line) or RU486-supplemented media (induced, solid line; *Myc* OE). No significant difference in survival between groups. Logrank (Mantel–Cox) test, n = 120 flies per condition across 3 cohorts, P > 0.05. **E**. Fitness assay (climbing performance) of flies containing both *Hml(Δ)-GeneSwitch* and *UAS-Myc.Z* maintained on control media (dashed line) or RU486-supplemented media (induced, solid line; *Myc* OE). Median climbing speed of flies during startle-induced negative geotaxis assays at indicated time points, measured using TrackMate in Fiji. Significant improvement in climbing performance was observed at days 49 and 56. Data presented as mean ± SEM, *n* = 30 flies per condition at study initiation (see Supplementary Table 5 for sample sizes at each timepoint), Mixed-effects model with Šídák’s multiple comparisons test, * Adj.P < 0.05. Genotypes are as follows: *w^1118^;srp-GAL4,UAS-GFP/+;crq-GAL4,UAS-GFP/+* (control, **A-C**), *w^1118^;srp-GAL4,UAS-GFP/UAS-Myc.Z;crq-GAL4,UAS-GFP/+* (*Myc* OE, **A-C**), *w* P{EP}Usf^G717^/w^1118^;srp-GAL4,UAS-GFP/+;crq-GAL4,UAS-GFP/+* (*Usf* OE, **A-C**), *w^1118^;UAS-Myc.Z/+; Hml(Δ)-GeneSwitch/+* (**D-E**).

